# RNA splicing programs define tissue compartments and cell types at single cell resolution

**DOI:** 10.1101/2021.05.01.442281

**Authors:** Julia Eve Olivieri, Roozbeh Dehghannasiri, Peter Wang, SoRi Jang, Antoine de Morree, Serena Y. Tan, Jingsi Ming, Angela Ruohao Wu, Tabula Sapiens Consortium, Stephen R. Quake, Mark A. Krasnow, Julia Salzman

**Author notes:** These authors contributed equally to this work (Julia Eve Olivieri, Roozbeh Dehghannasiri).

## Abstract

More than 95% of human genes are alternatively spliced. Yet, the extent splicing is regulated at single-cell resolution has remained controversial due to both available data and methods to interpret it. We apply the SpliZ, a new statistical approach that is agnostic to transcript annotation, to detect cell-type-specific regulated splicing in > 110K carefully annotated single cells from 12 human tissues. Using 10x data for discovery, 9.1% of genes with computable SpliZ scores are cell-type specifically spliced. These results are validated with RNA FISH, single cell PCR, and in high throughput with Smart-seq2. Regulated splicing is found in ubiquitously expressed genes such as actin light chain subunit *MYL6* and ribosomal protein *RPS24*, which has an epithelial-specific microexon. 13% of the statistically most variable splice sites in cell-type specifically regulated genes are also most variable in mouse lemur or mouse. SpliZ analysis further reveals 170 genes with regulated splicing during sperm development using, 10 of which are conserved in mouse and mouse lemur. The statistical properties of the SpliZ allow model-based identification of subpopulations within otherwise indistinguishable cells based on gene expression, illustrated by subpopulations of classical monocytes with stereotyped splicing, including an un-annotated exon, in *SAT1*, a Diamine acetyltransferase. Together, this unsupervised and annotation-free analysis of differential splicing in ultra high throughput droplet-based sequencing of human cells across multiple organs establishes splicing is regulated cell-type-specifically independent of gene expression.

## Introduction

Isoform-specific RNA expression is conserved in higher eukaryotes^1^, tissue-specific, and controls developmental^2–5^ and myriad cell signalling pathways^6,7^. Alternative splicing also plays a major functional role as it expands proteomic complexity and rewires protein interaction networks^8,9^. Alternative RNA isoforms of the same gene can even be translated into proteins with opposite functions^10^. Splicing is dysregulated in many diseases from neurological disorders to cancers^11^. Alternative splicing studies have been mostly limited to bulk-level analysis and they have shown evidence that as many as one third of all human genes express tissue-dependent dominant isoforms, while most of the highly-expressed human genes express a single dominant isoform in different tissues^12,13^. It has been known for decades that genes can have cell-type-specific splicing patterns, best characterized in the immune, muscle, and nervous systems^6,14–^16, 17^^. But, the extent of cell-type specific splicing is still controversial partly because it has only been studied indirectly through profiling tissues, which is confounded by differential cell-type composition. Many other questions remain such as whether cells of the same type in different tissues have shared splicing programs.

Determining how splicing is regulated in single cells could improve predictive models of splice isoform expression and move toward systems-level prediction of function. Furthermore, single-cell RNA splicing analysis has tremendous implications for biomedicine. Drugs targeting “genes’’ may actually target only a subset of isoforms of the gene and it is critically important to know which cells express these isoforms to predict on- and off-target drug interactions.

Genome-wide characterization of cell-type-specific splicing is still lacking mainly due to inherent challenges in scRNA-seq such as data sparsity. The field still debates whether single cell splicing heterogeneity constitutes another layer of splicing regulation or is dominated by stochastic splicing but stereotyped “binary” exon inclusion^18^, and whether cells’ spliced RNA is sequenced deeply enough in scRNA-seq for biologically meaningful inference. Most differential splicing analysis requires isoform estimation, which is unreliable with low or biased counts, or “percent spliced in” (PSI) point estimates, which suffer from high variance at low read depth and amplify the multiple hypothesis testing problem. Most methods for splicing analysis from scRNA-seq data are not designed for droplet-based data^19,20^. Studies of splicing in scRNA-seq data have focused on just a single cell type or organ, use pseudo-bulked data before differential splicing is analyzed, thus do not provide the potential to discover new subclusters or provide bona-fide quantification of splicing at single-cell resolution. Further, studies have almost exclusively used full-length data such as Smart-seq2^21,22,23^. Without genome-wide resolution, global splicing trends are missed and the focus on full-length sequencing data means that single cells sequenced with droplet-based technology, the majority of sequenced single cells including many cell types that are not captured by Smart-seq2^24,25^, have been neglected.

To overcome statistical challenges that have prevented analysis of cell-type-specific alternative splicing, we used the SpliZ^26^, a statistical approach that generalizes “Percent Spliced In” (PSI) (Figure 1A). For each gene, the SpliZ quantifies splicing deviation in each cell from the population average When cell type annotations are available, statistically identifies genes with cell-type-specific splicing patterns. The SpliZ is an unbiased and annotation-free algorithm and is applicable to both droplet-based and full-length scRNA-seq technologies. The SpliZ attains higher power to detect differential alternative splicing in scRNA-seq when genes are variably and sparsely sampled (Figure 1B) by controlling for sparsely-sampled counts and technological biases such as those introduced by 10x Chromium. Because the SpliZ gives a single value for each gene and each cell, it enables analyses beyond differential splicing between cell types, including correlation of splicing changes with developmental trajectories and subcluster discovery within cell types based on splicing differences (Figure 1C-D). It also provides a statistical, completely annotation-free approach that identifies splice sites called SpliZsites that contribute most variation to cell-type specific splicing.

**Figure 1.**
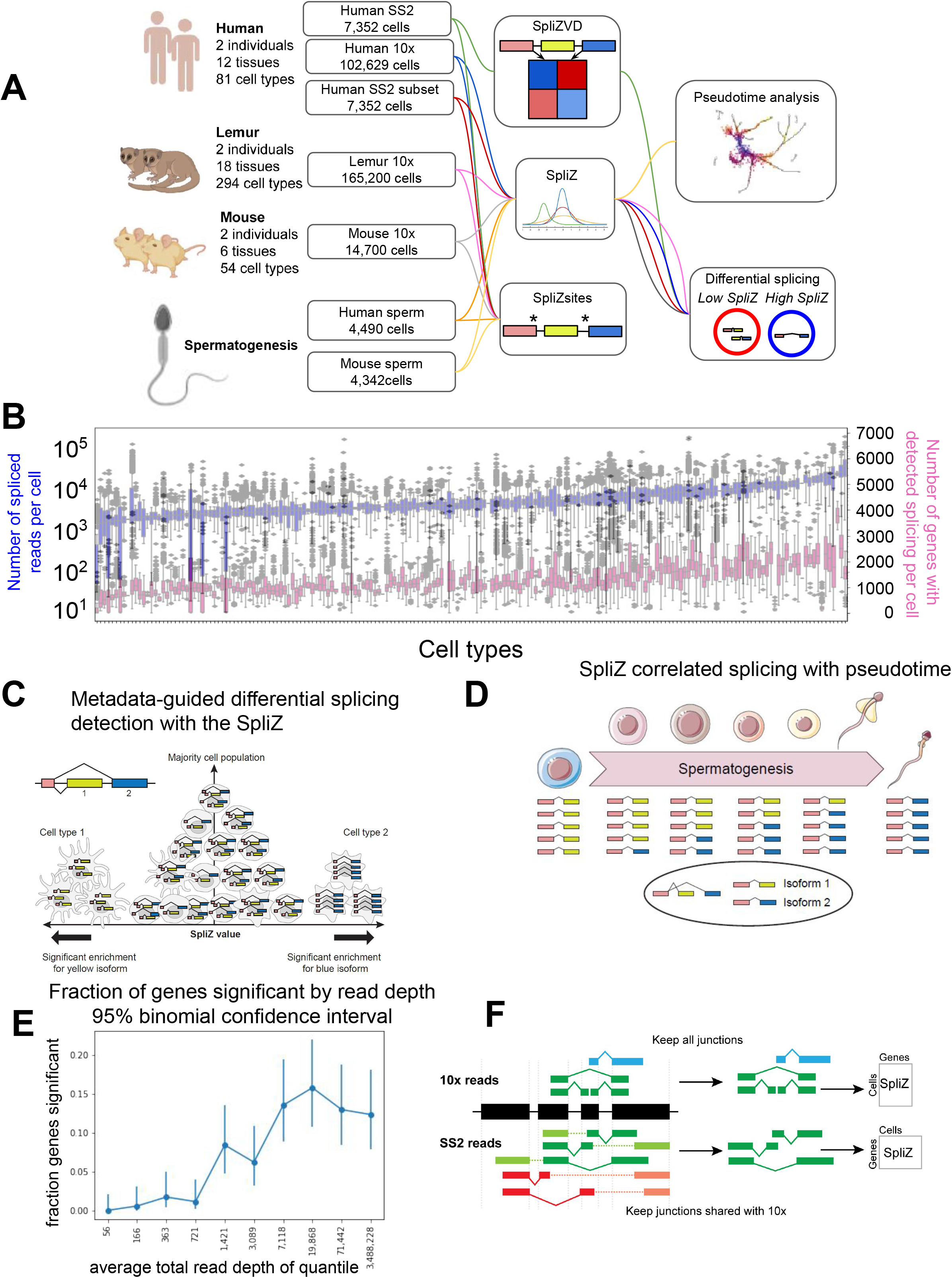
Analysis of alternative splicing in single cell RNA-seq. (A) Human, mouse lemur, and mouse single cell RNA-seq from 10x and Smart-seq2 were run through the SpliZ pipeline for differential splicing discovery. (B) 10x data from the first human individual contains 82 cell types with variable sequencing depth. (C) Given cell type annotation, SpliZ scores can be aggregated for each cell type, allowing identification of cell types with statistically deviant splicing. (D) Cell-wise SpliZ values can be correlated with pseudotime. (E) The fraction of genes called as having significant differential alternative splicing by cell type is higher at higher sequencing depths, plateauing at around 20,000 spliced reads in the dataset, at which point around 15% of genes were called as significant. (F) The SpliZ is calculated independently for Smart-seq2 data restricted to junctions found in 10x data, and used to validate results from 10x data.

Here, we used the SpliZ to analyze 75,146 cells profiled with 10x Chromium (10x) across 12 tissues and 82 cell types from one human individual through the *Tabula Sapiens* project^27^. We also performed SpliZ analysis on a second human and two mouse lemur and two mouse individuals: together the analysis is applied to 117,333 human^27^, 165,200 mouse lemur^28^, and 14,700 mouse cells^29^ with 10x. Additionally, we analyzed a developmental trajectory during spermatogenesis course 4,490, 4,342, and 5,601 sperm cells from human, mouse, and mouse lemur^28^, respectively. The SpliZ has higher power to detect differential alternative splicing between cell types at higher sequencing depths, plateauing at around 20,000 spliced reads measured for the gene, at which point around 15% of genes were called as significant (Figure 1E).

We performed high-throughput computational validation with the Smart-seq2 cells and the data from the along with experimental and in-situ validations including Sanger sequencing and RNA FISH (Figure 1F). Mouse and mouse lemur data was used to assess evolutionary conservation of the discoveries in the human. Examples found by this analysis include differential cell-type-specific and compartment-specific alternative splicing in a small subset of ubiquitously expressed genes including *MYL6*, an actin light chain subunit, *RPS24*, a core ribosomal subunit associated with Diamond-Blackfan Anemia^22^, and *TPM1*, a tumor suppressor tropomyosin. Furthermore, we identify regulated splicing changes during spermatogenesis, including new findings of conserved regulated splicing in genes *CEP112* and *SPTY2D1OS*.

To our knowledge, this work provides the first unbiased and systematic screen for cell-type-specific splicing regulation in highly resolved human cells, predicting functionally significant alternative splicing and calls for more attention to the potential of scRNA-seq for discovering regulated splicing in single cells.

## RESULTS

### Conserved splicing in ubiquitously expressed genes, including *ATP5FC1* and *RPS24*, predicts cellular compartment at single cell resolution

We applied the SpliZ^26^, a recently-developed method to identify cell-type-specific splicing to ∼75k 10x cells in 12 tissues from one human donor^27^, beginning by testing for splicing regulation differing by tissue compartment (immune, epithelial, endothelial, and stromal) regardless of the tissue of origin. This analysis identified 1.6% (22 of 1,353) of genes with calculable SpliZ scores as having consistent compartment-specific splicing effects (Methods). *ATP5F1C, RPS24*, and *MYL6*, have the highest compartment-specific splicing effects, defined as the largest magnitude median SpliZ in any compartment (Supp. Figure 1). Their compartment-specific splicing was conserved in mouse and mouse lemur. *ATP5F1C* is the gamma subunit of mitochondrial ATP synthase, a multi-subunit molecular motor that converts the energy of the proton potential across the mitochondrial membrane into ATP. *MYL6* is an actin light chain subunit known to have cell-type-specific splicing differences in the muscle^30^. *RPS24* is an essential ribosomal protein for ribosome small subunit 40S discussed in detail later. Among the examples of genes demonstrating compartment-specific splicing is *LIMCH1* (Figure 6B). *LIMCH1* has been reported as a non-muscle myosin regulator^31^ and has been associated with Huntington’s disease^32^ with little other characterization, including, to our knowledge, no reports of regulated splicing.

To test the predictive power of compartment-specific genes at single cell resolution, we performed unsupervised k-means clustering on the SpliZ scores of *RPS24* and *ATP5F1C* alone. Setting k=3, cells from stromal, epithelial, and immune compartments were classified with accuracies of 78%, 84%, and 95% respectively independent of gene expression (Figure 2, Methods). This establishes that splicing of a minimal set of genes, in this case only 2, has high predictive power of the compartmental origin of each single cell. Underlining tight biological regulation of splicing in these genes, parallel analysis in the 10x scRNA-seq data from mouse lemur and mouse shows compartment-specific splicing is conserved for RPS24 and MYL6 (Figures 3-4).

**Figure 2.**
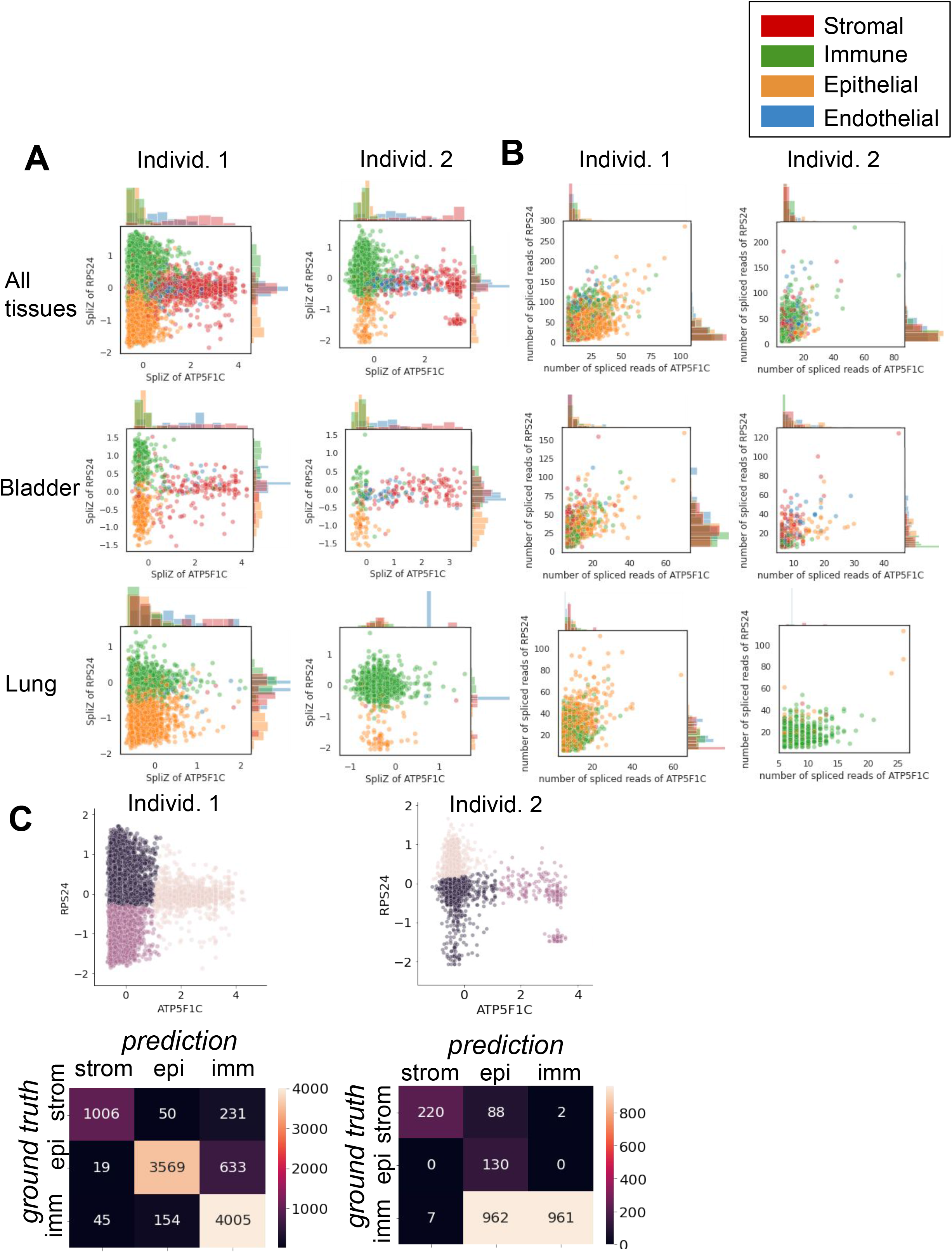
(A) The SpliZ scores of the genes *ATP5F1C* and *RPS24* together separate compartments in both human individuals. (B) Using the spliced read counts for each gene rather than the SpliZ does not separate the compartments, showing that this separation is not driven by gene expression differences. (C) Unsupervised k means clustering results in 78%, 84%, and 95% accuracy of compartment classification for the stromal, epithelial, and immune compartments respectively for individual 1, and 70%, 100%, and 49% respectively for individual 2.

**Figure 3.**
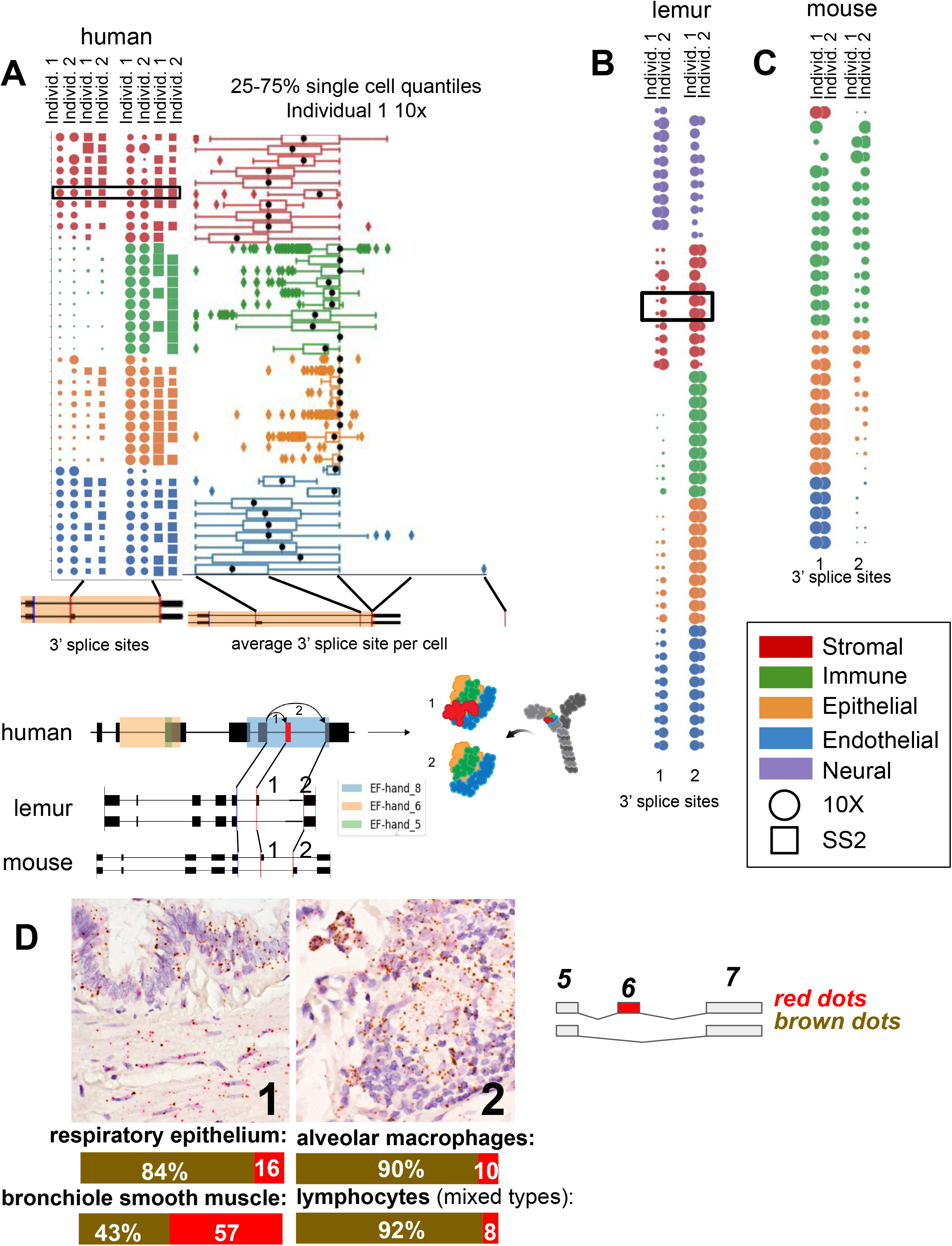
Differential alternative splicing between compartments for *MYL6* is driven by an exon skipping event with orthologous splice sites in (A) human, (B) lemur, and (C) mouse. Each dot plot shows junction expression for the splice site marked in blue on the gene annotation, with dot size proportional to the fraction of junctional reads from the given blue splice site (marked on the gene annotation) to each of the splice sites marked in red (numbered on the x axis of the dot plots). Dots are colored by compartment. Columns of dots are biological replicates; for human data the first two columns are 10x samples (circles) and the second two columns are Smart-seq2 samples (squares). For mouse and lemur the two columns are 10x samples. Colored blocks on the gene annotation identify known protein domains. Cells in the immune compartment have higher exon skipping rates than cells in the stromal compartment in all three organisms. Smooth muscle cell types are boxed within the stromal compartment. Mouse cells have the same relative proportions of exon inclusion between compartments, but express higher levels of the exon included isoform overall. (D) RNA FISH validation in human lung: Each slide is stained simultaneously with probes in red (specific to exon inclusion) and brown (specific to exon exclusion). As found from the scRNA-seq data, smooth muscle cells have a higher proportion of the included exon than the other compartments and immune cells have the lowest proportion.

### The splicing of actin regulator *MYL6* is compartment-specifically regulated

We identified *MYL6* as both cell-type-specifically and compartment-specifically spliced in humans and its splicing pattern is conserved. *MYL6* is a ubiquitously expressed myosin light chain subunit and is known to have a lower level of exon skipping in muscle than non-muscle tissue^30^, but differential exon skipping at a single-cell level has only been characterized in smooth muscle cells. We find in human, mouse, and mouse lemur, the stromal compartment, which includes smooth muscle, as well as the endothelial compartment have a relatively higher proportion of exon inclusion, while the epithelial compartment has a lower level of exon inclusion and the immune compartment has the lowest level of exon inclusion (Figures 3A-C, Supp. Figure 2).

We validated compartment-specific differential alternative splicing in *MYL6* using RNA FISH with isoform-specific probes on human adult lung tissue obtained from the Stanford Tissue Bank (Figure 3D, Methods). In human lung, this confirmed that bronchiole smooth muscle cells have the highest fraction of the exon retention isoform (57%), while the respiratory epithelium has a lower fraction of the +exon isoform (16%) and the two immune cell types profiled have the lowest fractions of the +exon isoform (10% and 8%). We separately performed RNA FISH on human cells isolated from the muscle which showed that mesenchymal stem cells and muscle stem cells have a higher proportion of the +exon isoform than endothelial cells (Supp. Figure 3A, Methods).

### *RPS24* has compartmentally-regulated alternative splicing and expresses a microexon in human epithelial cells

*RPS24* is a highly expressed and essential ribosomal protein. Our analysis revealed that *RPS24* has the most significant cell-type-specific and compartment-specific alternative splicing patterns at its C terminus in human and mouse lemur. The significance of the alternative splicing patterns of *RPS24* is underscored by recent findings that ribosome composition is more modular than previously appreciated in a cell and tissue specific manner^33^. There has been a partial study of *RPS24* splicing treating two isoforms^19^, and another study reported modest differential splicing at the tissue level^19,22^ for *RPS24* involving three isoforms. However, here, we show that splicing of *RPS24* is complex, highly regulated at a single cell level.

Differential alternative splicing of RPS24 involves alternate inclusion of three short exons, a, b, and c, each only 3, 18, and 22 nucleotides long, respectively (Figure 4A-B); regions of genomic sequence around exon a is ultraconserved. Splicing of a,b,c exons results in isoforms of the protein that differ by the presence of a single lysine at the solvent exposed site of the ribosome prompt a hypothesis that they affect post-translational modifications ^34^. The -a-b+c isoform is dominant in all endothelial cell types in human, as well as most stromal cell types and half of the immune cell types. Within the immune compartment, our global analysis reveals differential alternative splicing of *RPS24* in monocytes, where the -a-b-c isoform is dominant in classical monocytes residing in multiple tissues and the -a-b+c isoform is dominant in non-classical monocytes. Intermediate monocytes have equal proportions of each.

**Figure 4.**
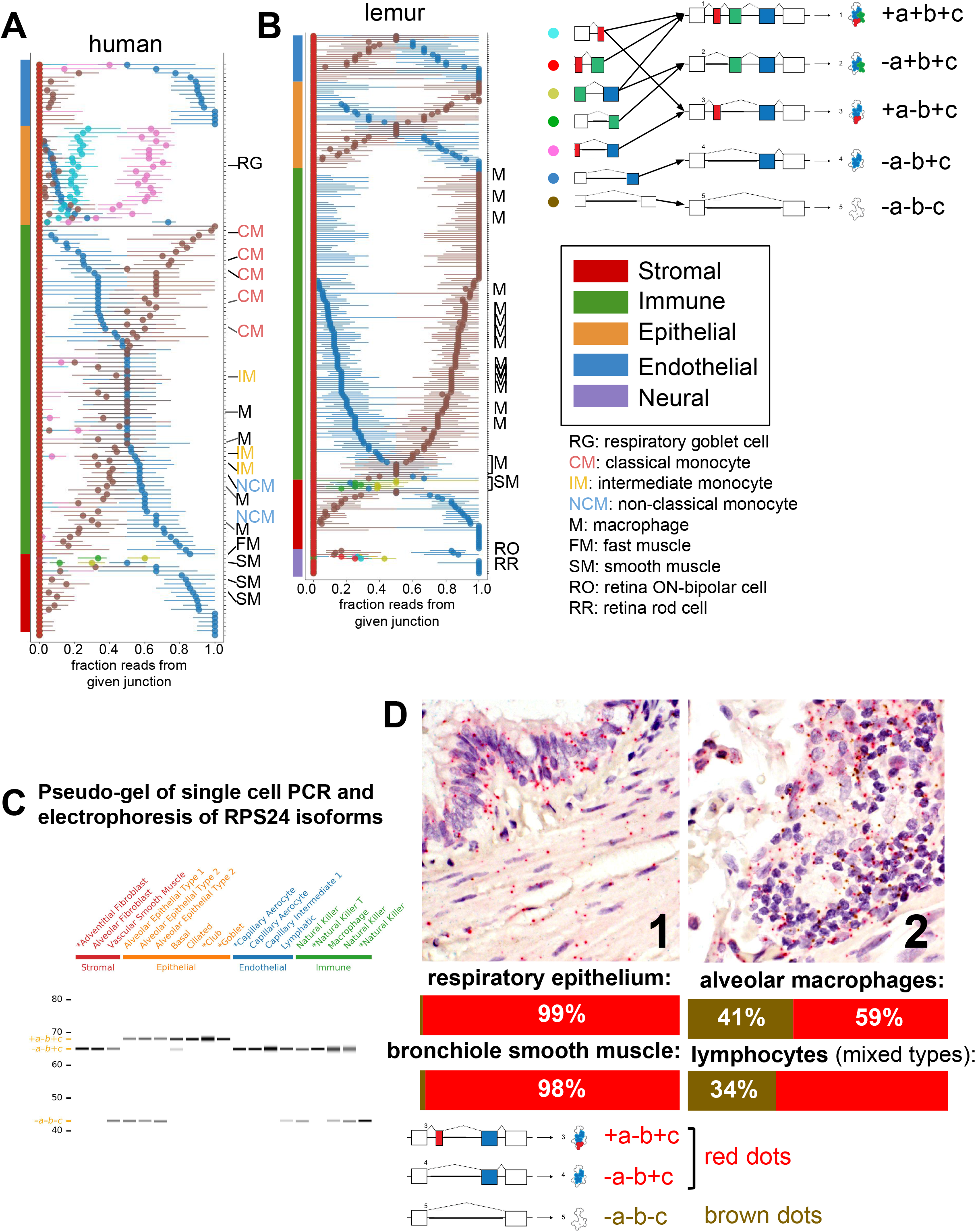
(A) *RPS24* has striking compartment-specific alternative splicing. Each color in the plot represents one *RPS24* junction that uniquely identifies an isoform (green represents two isoforms). For each cell type (y axis), the median of all single-cell point estimates of junction fraction in the cell type is plotted on the x axis, with bars representing the 25th and 75th quantiles of single-cell junction fractions. Cell types are sorted by compartment. Gene annotation corresponds to isoform and impact on *RPS24* protein. Within the immune compartment, the fraction of the blue junction increases from classical monocytes to intermediate monocytes to non-classical monocytes. (B) The isoform with epithelial-specific splicing in human is not expressed in lemur. However, the same isoform is expressed in smooth muscle as in human. Retinal cells are the only cells to express the +a+b+c isoform. (C) Single-cell PCR validates prediction that the +a-b+c isoform is epithelial-specific. All the epithelial cells contain the isoform with the 3-nt exon 5a, while none of the cells from other compartments do. PCR products from the cells prefixed by asterisks were Sanger-sequenced and matched the expected isoform without evidence of mixture. (D) *RPS24* FISH in human lung validates scRNA-seq predictions. Slides simultaneously stained with probes in red and brown, specific for alternative splice junctions. As found from the scRNA-seq data, respiratory epithelium and bronchiole smooth muscle, in the epithelial and stromal compartments respectively, have a low proportion of the -a-b-c isoform compared to alveolar macrophages and lymphocytes, both of which are in the immune compartment.

Epithelial cell types in human are marked by the dominance of the +a-b+c isoform, which is barely present in any non-epithelial cell types and is not dominant in any of them. The +a-b+c isoform is only found at very low levels in mouse and mouse lemur (Supplement). The human epithelial specificity of the +a-b+c isoform is further supported by single-cell RT-PCR (Figure 4C).

Other cell types have distinct isoform expression: the -a+b+c isoform is specific to fast and smooth muscle cells in both human and lemur, such as thymus fast muscle and bladder smooth muscle in human, as well as vascular associated bladder smooth muscle in mouse lemur, though some smooth muscle cell types in human do not express it, specifically thymus vascular associated smooth muscle and vasculature smooth muscle. Among profiled cell types, +a+b+c isoform is found only in retinal cells in the mouse lemur (retina data not available for human).

In addition to high throughput validation using bowtie2 alignment (Supp.Figure 4), we performed RNA FISH to independently validate cell-type-specific splicing in a subset of lung and muscle cells (Figure 4D, Supp. Figure 3B). This data confirms that the -a-b-c isoform composes just ∼1-2% in the respiratory epithelium and bronchiole smooth muscle, while alveolar macrophages and lymphocytes have about 34-41% -a-b-c.

### ∼9% of measured genes have cell-type-specific splicing regulation

Splicing regulation in the vast majority of human cell types has not been characterized. We used the 82 annotated cell types in Tabula Sapiens to identify genes with statistical support for having differential alternative splicing patterns using the same procedure applied to identify compartment-specific genes (Figure 1C, Methods)^26^. 129 out of 1416 genes (9%) had significant cell-type-specific splicing profiles based on discovery with 10x data (p value < 0.05, effect size > 0.5) (Methods, Supp. Figure 5). Among genes called significant, the Pearson correlation between median SpliZ in individuals 1 and 2 (10x) was 0.77 and 0.44 (p<10e-50) between 10x and Smart-seq2 within individual 1 (Methods, Supp. Figure 6).

Tropomyosin 1, *TPM1* which has three isoforms each impacting the tropomyosin domain at the 3’ end of the transcript, served as a positive control in this analysis as it is known to undergo cell-type specific splicing; it is ranked as the 27th most significant effect size. Unbiased SplIZ analysis finds that capillary endothelial cells express about equal levels of three isoforms, smooth muscle cells almost exclusively express the isoform with the 3’-most domain. (Figure 5A-C). This trend, among others, is consistent with knowledge of *TPM1* splicing from other studies^35^, though extending its splicing profiles to cell types where it has never been characterized. Among the comprehensive catalog of differences, mesothelial cells and pericyte cells consistently include different cassette exons at the 3’ end of the transcript, affecting the tropomyosin protein domain. Splicing biology *TPM2* and *TPM3*, two other genes from the TPM family where partial characterization has suggested cell-type specific splicing are similarly rediscovered in our analysis but significantly extended: cell types outside of muscle such as bladder pericytes and bladder fibroblasts have similar splicing profiles to smooth or fast muscle cells respectively (Figure 5).

**Figure 5.**
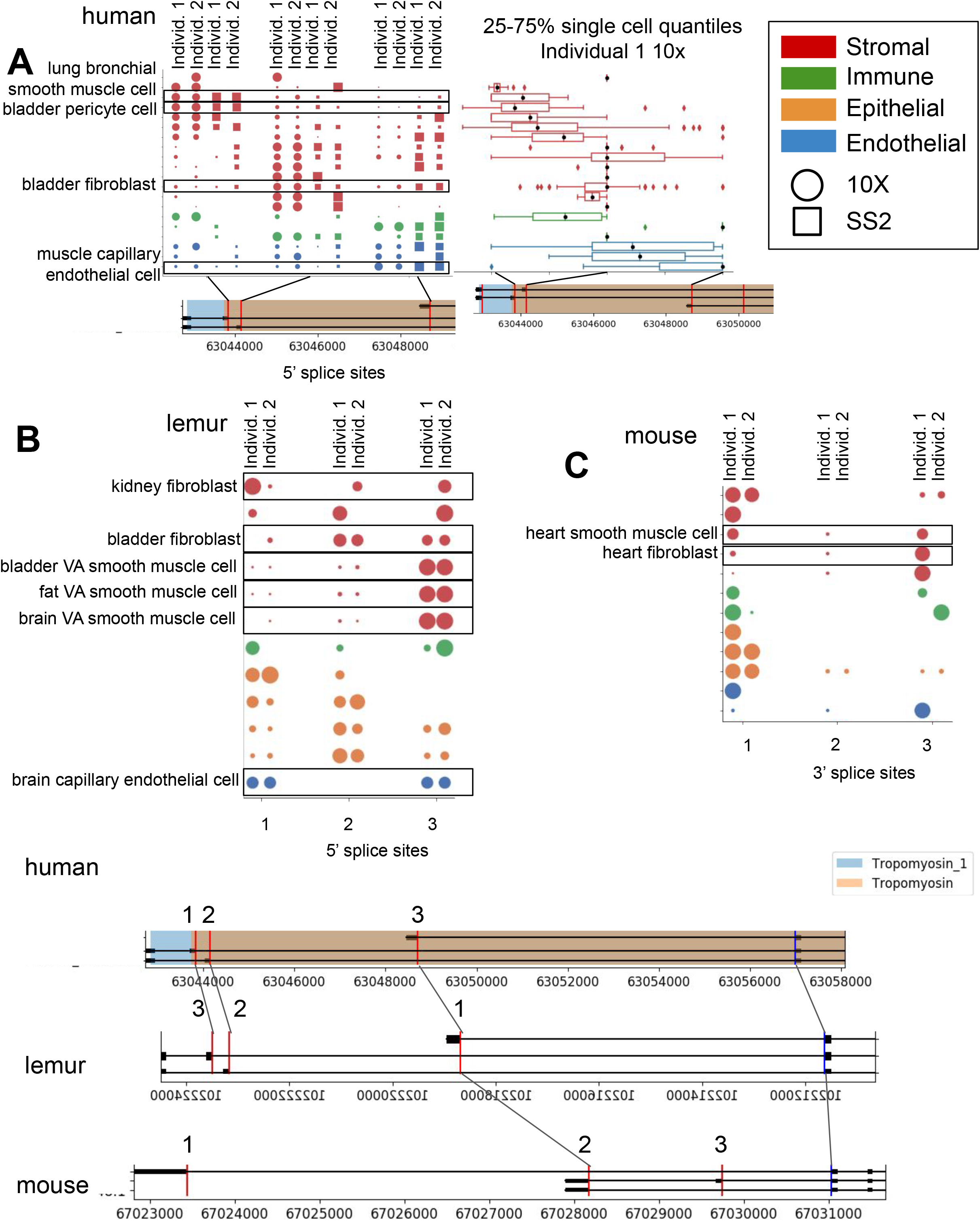
Conserved splicing in *TPM1* is recovered in (A) human, (B) mouse lemur, and (C) mouse. *TPM1* has a pattern of differential splicing involving two cassette exons and an alternate 3’ end. Capillary endothelial cells mostly express the isoform with the alternate 3’ end, while smooth muscle almost exclusively expresses the isoform with the 3’-most domain (b0xed in the figure). There is differential isoform usage within the stromal compartment as well, for example bladder stromal fibroblasts and bladder stromal pericytes each express a different dominant cassette exon. Both lemur and mouse similarly express cell-type specific differences in *TPM1* isoform usage. See Figure 3 for explanation of plots.

*PNRC1*, a nuclear receptor coregulator that functions as a tumor suppressor as the 5th highest effect size (Figure 6C), ^36^; limited in vitro studies have found evidence that splice variants of *PNRC1* modify its interaction domains and nuclear functions^37^. The largest magnitude median SpliZ score is found in muscle stromal mesenchymal stem cells, revealing new biology. Other cell types, including immune and stromal types in the bladder, have markedly distinct splicing programs (Figure 6).

**Figure 6.**
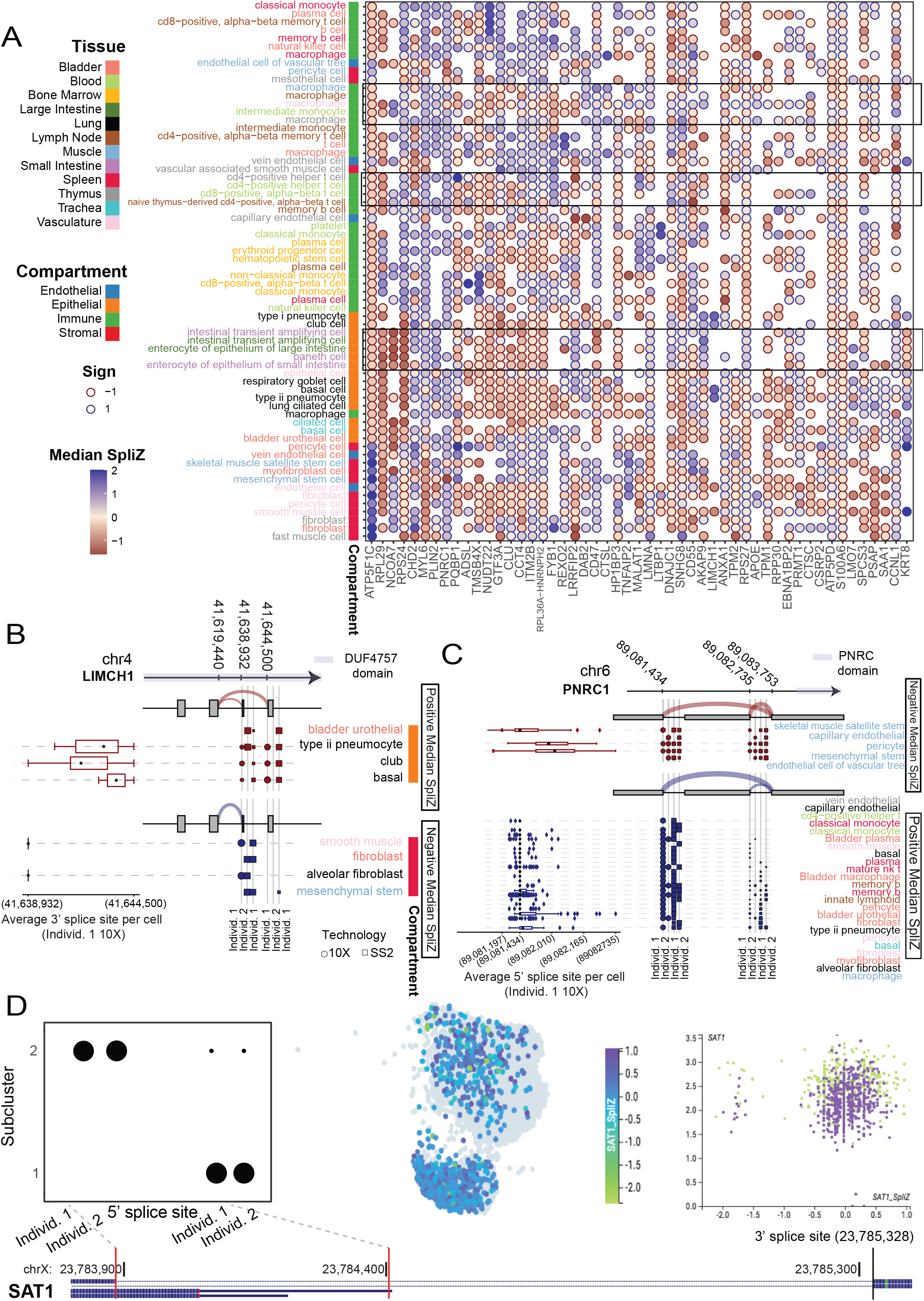
(A) Unsupervised clustering analysis with the SpliZ identified clusters of cell types and compartments independent of tissue. Dots show the median SpliZ (effect size) for genes found to be significantly regulated across cell types. Only 50 significant genes with the highest effect size and cell types with > 25 significant genes are shown. Hierarchical clustering was performed on both genes and cell types based on median SpliZ values. Cell type names are color-coded based on their tissue and the sidebar shows the compartment for each cell type. (B) Dot and Sashimi plots showing *LIMCH1* differential exon skipping impacting a protein domain of unknown function DUF4757 across cell types and human datasets. Vasculature smooth muscle cells and fibroblasts always include the exon (at 41,638,932) while bladder urothelial cells skip with >50% rate. Cell types are colored according to their tissues of origin and the same tissue and compartment colors are used as in (A). (C) Differential exon skipping by cell type in *PNRC1* is replicated across the four human datasets. Skeletal muscle satellite stem cells include the exon about 50% of the time, whereas vein endothelial cells in the thymus never include the exon. Cell types with negative median SpliZ values are on the top panel and those with positive median SpliZ values are on the bottom panel. Each of the four human datasets is plotted, with circles representing 10x data and squares representing Smart-Seq2 data. The gene annotation is shown above the dots, with arcs indicating the median expression of each junction for the given panel. Known protein domains are marked on the gene structure. Box plots for each cell type based on the SpliZ for each cell in the cell type in individual 1 10x data (L). Each box shows 25-75% quantiles of average donor per cell. (D) Alternative splicing of gene *SAT1* distinguishes two populations of cells within blood classical monocyte cell type. The alternative splicing involves one annotated and one unannotated splice junction in gene *SAT1*. Dot plot for the differential inclusions of the 5’ splice sites for the 3’ splice site at 23,785,300 for the cells grouped based on the subclusters obtained via model-based Gaussian mixture model clustering. Clustering based on gene expression (as shown by cellxgene visualization) does not distinguish cell populations with distinct splice profiles. Visualization of the Gene expression value for *SAT1* does not distinguish the populations; both subclusters contain classical monocytes from both human individuals (right scatter plot). The number of reads (X) and cells (Y) containing the splice junctions involving the 3’ splice site in each individual at right

The high dimensionality of SpliZ scores enabled us to test if unsupervised clustering on the median SpliZ scores could recapitulate relationships between cell types. For example, many immune cell types and muscle cell types are detected in multiple tissues. We found that the same cell types from different tissues are generally clustered together, including macrophages and T cells from different tissues and also intestinal cell types from large and small intestine (Figure 6A). This clustering also reveals that the splicing programs of cell types from the same compartment are highly similar and automatically clustered together independent of their tissues (Figure 6A).

### The most statistically variable splice sites with cell-type-specific regulated splicing are annotated splice sites involved in unannotated alternative splicing

The biological importance of splicing detected by the SpliZ and that it is completely agnostic to isoform annotation led us to test whether cell-type specific splicing variation is (a) focused at exons that are annotated to undergo alternative splicing and (b) conserved. The SpliZ method uses a statistical, annotation-free approach to identify SpliZsites: variable splice sites that contribute most to the cell-type specific splicing of a gene agnostic to gene annotation. SpliZsites blindly re-identify known alternative splice sites in *ATP5F1C, MYL6, TPM1*, and *RPS24*. In *TPM2*, the SpliZ re-identifies two known alternative splicing sites but also predicts a cell-type-specific un-annotated alternative splicing event in stromal cell types involving 5’ splice site 35684315 affecting Tropomyosin and Tropomyosin_1 protein domains.

Genome-wide, the vast majority of SpliZsites (92%) in significant genes are at boundaries of annotated exons. However, only 32% were annotated as involved in alternatively spliced isoforms, suggesting that un-annotated-- rather than annotated-- exon skipping accounts for underappreciated splicing variation in single cells. Supporting the idea that SpliZsites discover real biological signals, 13% of the SpliZsites in the human are also the SpliZsites in the mouse lemur and/or mouse (Methods). Further, exon skipping has a global effect on single-cell proteomes: more than half of SpliZsites impacted protein coding domains; 25% impact the 3’UTR and 16% impact the 5’ UTR, consistent with a bias in 10x technology towards the 3’ gene end.

### The SpliZ identifies subpopulations of classical monocytes with distinct splicing programs in SAT1

The SpliZ has a theoretical normal distribution under the assumption that cells within a cell type all have the same propensity to express each splice isoform (the “null hypothesis”). This property allows us to statistically test whether cell types subcluster on the basis of splice isoform, as quantified by the SpliZ using an integrated completed likelihood (ICL) model selection framework based on Gaussian mixture modeling. Importantly, this method avoids false positives of apparent binary exon inclusion^18^(Methods, manuscript in preparation). We applied this method to immune cell types to illustrate the power of single-cell splicing quantification by the SpliZ for defining subsets of cells within an annotated cell types defined by gene expression. Methods that detect splicing changes based on pseudo bulk rely on metadata and could not identify subsets of cells within an annotated cell type.

Analysis of classical monocytes based on the SpliZ discovered distinct populations of cells with distinct splice profiles in *SAT1*, the gene with most statistical evidence of distinct splicing in two clusters in our analysis. Cells were subset according to cluster assignment based on Gaussian-mixture-model-based clustering (Methods). SpliZsites driving this separation show a clear pattern of distinct isoform expression profiles: cells in cluster 1 splice to an unannotated 5’ splice site, whereas those in cluster 2 contain an annotated splice junction. We then blindly tested whether classical monocytes in individual 2 exhibited the same two clusters of distinct splicing programs; they did (Figure 6D). These clusters are not driven by *SAT1* gene expression and are not detected by gene-based clustering of monocytes as shown by visualization in cellxgene^38^ (Figure 6D). Further supporting a biological role of *SAT1* splicing in the immune system, the same populations of *SAT1* splicing profiles exist in lung and thymus. Together with statistical support, and blinded validation, this supports that *SAT1* exhibits splicing programs that define two cell states within classical monocytes and calls for further investigations for up and downstream regulation. Other genes including *PTP4A2, RABAC1*, and *MAGOH* have similar evidence of sub-population structure and warrant further investigation.

### The SpliZ discovers conserved alternative splicing in mammalian spermatogenesis

SpliZ provides a systematic framework to discover how splicing is regulated at a single cell level along developmental trajectories (i.e., pseudotime). Previous studies have shown that testis is among the tissues with the highest isoform complexity and that even the isoforms diversity in spermatogenic cells (spermatogonia, spermatocytes, round spermatids, and spermatozoa) is higher than that of many tissues^39^. Also, RNA processing has been known to be important in spermatogenesis^40^. However, alternative splicing in single cells during spermatogenesis at the resolution of developmental time enabled by single cell trajectory inference has not been studied. To systematically identify cells whose splicing is regulated during spermatogenesis, we applied SpliZ to 4,490 human sperm cells^41^ and compared findings to mouse^41^ and mouse lemur ^28^ sperm cells to test for conservation of regulated splicing changes.

170 genes out of 1,757 genes with calculable SpliZ in >100 human cells have splicing patterns that are significantly correlated with the pseudotime (|Spearman’s correlation| >0.1, Bonferroni-corrected p-value < 0.05). The highest correlated genes included *TPPP2*, a gene regulating tubulin polymerization implicated in male infertility^42^, *FAM71E1*, being predominantly expressed in testes^43^, *SPATA42* a long non-coding RNA implicated in azoospermia^44^, *MTFR1*, a gene regulating mitochondrial fission, and *MLF1*, an oncogene regulated in drosophila testes^45^(Suppl. Figure). In *MTFR1*, SpliZsites identify an unannotated 3’ splice site in immature sperm showing evidence of novel transcripts (Figure 7A). The regulated splicing programs in *TPPP2* involves a highly expressed un-annotated splice isoform that modifies the functionally annotated p25-alpha domain thought to be brain-specifically expressed^46^ (Figures 7C).

**Figure 7.**
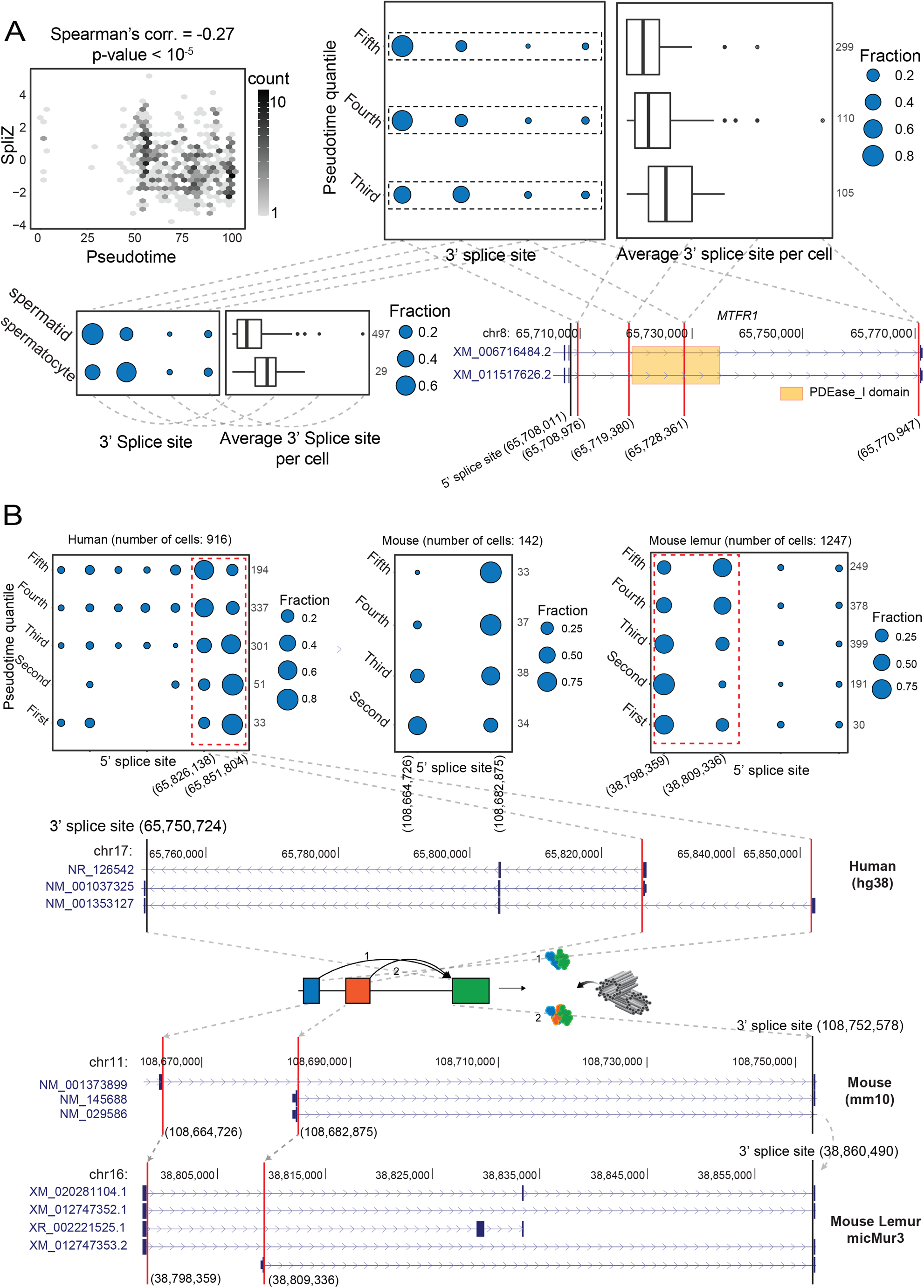
(A) Regulated alternative splicing of *MTFR1* during sperm development. Significant negative correlation (Spearman’s correlation = -0.27, p-value =1.23e-7) between the SpliZ score for gene *MTFR1* and the pseudotime in human sperm cells (top left). Dot plot and box plot show increasing use of a downstream 3’ Splice site driving the *MTFR1* SpliZ in equal pseudotime quantiles (top right) with the same trend in immature (spermatocyte) and mature (spermatid) cells (bottom left). The gene structure for *MTFR1* according to human RefSeq annotation database is shown (bottom right). (B) Dot plots showing the developmentally-regulated alternative splicing of gene *CEP112* in testis cells from human, mouse, and mouse lemur. Cells grouped according to pseudotime quantiles. The alternative splicing is conserved (i.e., involves the same set of 5’ and 3’ splice sites in human, mouse, and mouse lemur data) and involves 5’ splice sites 65,826,138 and 65,851,804 in human, 5’ splice sites 108,664726 and 108,682,875 in mouse, and 5’ splice sites 38,798,359 and 38,809,336 in mouse lemur. *CEP112* is on the plus strand in the human genome but is on the minus strand in mouse and mouse lemur genomes. Gray arrows show the liftover mapping between the 3’ splice site and two 5’ splice sites of the exon skipping event and gray dashed lines for the human plot show the location of the 5’ splice sites and how splicing changes the Apolipoprotein protein domain.

Among significantly correlated genes in human cells, splicing in 10 of these genes is also developmentally regulated in mouse and mouse lemur. Centrosomal protein 112 (*CEP112*), a coiled-coil domain containing centrosomal protein and member of the cell division control protein had the highest SpliZ-pseudotime correlation. It is highly expressed in testes and is essential for maintaining sperm function: loss-of-function mutations in *CEP112* have recently been associated with male infertility^47^. Strikingly, the same 3’ splice site and 5’ splice sites identified by SpliZsites are orthologous and affect the apolipoprotein domain, a protein involved in the delivery of lipid between cell membranes an is critical for the sperm development and fertility^48^ (Figure 7D).

*SPTY2D1OS* is another gene with conserved regulated splicing in spermatogenesis (Suppl. Figure 8). Though highly expressed in human testes, it has unknown function in sperm development. *SPTY2D1OS* is located between *Uveld* and *SPTY2D* in the human genome; in mouse, *SPTY2D1OS* corresponds to a lncRNA named Sirena1, which has been recently shown to have function in mouse oocyte development^49^ but not previously implicated in spermatogenesis. Together our results suggest transcriptome-wide regulation of splicing in spermatogenic cells and calls for more investigation into the function of splicing regulation not only in sperm development but also in other developmental trajectories.

## Conclusion

Cell-type-specific splicing has been known to have functional effects in some cell types and in some genes for decades. However, technological limitations in measurement technology and methods to analyze resulting data have prevented high throughput studies that profile the extent to which cell type can be predicted from splicing information alone. Full-coverage technologies such as Smart-seq2 have been the primary technologies for analyzing splicing in single cells thus far. However, Smart-seq2 is very difficult to scale: sequencing of 5,000 cells that would take 2-3 days using 10x is estimated to take ∼26 weeks using Smart-seq2^24,50^. The lack of analysis of splicing in droplet data prevents discovery of regulated splicing in cell types that cannot be adequately profiled by plate-based approaches, and therefore causes biologically regulated splicing in these cell types to be missed^25^. Here, we apply new analytic methodologies to find highly regulated splicing patterns from ubiquitous droplet based sequencing platforms. These results reveal deeply conserved splicing programs that define tissue compartment and cell type in vivo.

The reproducibility of models in independently generated datasets suggest that the SpliZ can be applied globally to larger numbers of cell types to further identify splicing regulation at a single cell level. We predict that as the number of cells profiled and the cell types grow, and global analyses are performed on data that is not 3’ biased, the fraction of genes with evidence of cell-type-specific splicing will increase substantially beyond our current estimate of around 10% (Figure 1F). By providing single-cell resolved quantification, the SpliZ can also be used to compare closely related cell types across diverged organisms to generate functional hypotheses of which splicing programs are most conserved and which may be candidates that nominate evolutionary innovations. Smart-seq2 is The results presented here lay the foundation for comprehensive splicing analysis in any scRNA-seq dataset and a reference to which future studies can be compared. Together, this work provides strong evidence for the hypothesis that alternative splicing in a large majority of human genes is regulated in the majority of cell types and supports the idea that splicing is central to functional specialization of cell types.

## Acknowledgments

We thank Manny Ares, Douglas Black, Maria Barna and members of the *Tabula Sapiens* and Salzman lab for insightful discussions. We thank Jessica Klein for creating part of Figure 1, and also Emma Chory for sashimi plots. J.O. is supported by the National Science Foundation Graduate Research Fellowship under Grant No. DGE-1656518 and a Stanford Graduate Fellowship. R.D. is supported by the Cancer Systems Biology Scholars Program Grant R25 CA180993 and the Clinical Data Science Fellowship Grant T15 LM7033-36. J.S. is supported by the National Institute of General Medical Sciences Grant R01 GM116847 and NSF Faculty Early Career Development Program Award MCB1552196.

## METHODS

### File downloads

- Human RefSeq hg38 annotation file was downloaded from: ftp://ftp.ncbi.nlm.nih.gov/refseq/H_sapiens/annotation/GRCh38_latest/refseq_identifiers/GRCh38_latest_genomic.gff.gz
- Mouse Lemur RefSeq Micmur3 annotation file was downloaded from: https://www.ncbi.nlm.nih.gov/assembly/GCF_000165445.2/
- Mouse RefSeq GRCm38.p6 annotation file was downloaded from: https://ftp.ncbi.nlm.nih.gov/genomes/all/GCF/000/001/635/GCF_000001635.26_GRCm38.p6/GCF_000001635.26_GRCm38.p6_genomic.gtf.gz
- The UCSC Pfam database for the human hg38 genome assembly was downloaded from: http://hgdownload.soe.ucsc.edu/goldenPath/hg38/database/ucscGenePfam.txt.gz

### Code Availability

Code to reproduce analysis and create figures is available through this GitHub repository: https://github.com/juliaolivieri/DiffSplice.

### Data Availability

The fastq files for the Tabula Sapiens data (both 10x and Smart-seq2) were downloaded from https://tabula-sapiens-portal.ds.czbiohub.org/. The pilot 2 individual is referred to as individual 1, and the pilot 1 individual is referred to as individual 2 in this manuscript. Pancreas data was removed from individual 2. Cell type annotations were downloaded on March 19th, 2021, and the “ground truth” column was used as the within-tissue-compartment cell type. The Tabula Muris data was downloaded from a public AWS S3 bucket according to https://registry.opendata.aws/tabula-muris-senis/. The P1 mouse is referred to as individual 1 and P2 is referred to as individual 2 in this manuscript. Compartment annotations were assigned based on knowledge of cell type. The fastq files for the Tabula Microcebus mouse lemur data were downloaded from https://tabula-microcebus.ds.czbiohub.org. Antoine is referred to as individual 1 and Stumpy is referred to as individual 2 in this manuscript. The propagated_cell_ontology_class column was used as the within-tissue-compartment cell type. Because tissue compartments in the mouse lemur were annotated more finely, we collapsed the lymphoid, myeloid, and megakaryocyte-erythroid compartments into the immune compartment. Human and mouse unselected spermatogenesis data was downloaded from the SRA databases with accession IDs SRR6459190, SRR6459191, and SRR6459192 for human, and accession IDs SRR6459155, SRR6459156, and SRR6459157 for mouse.

### How SpliZ pipeline was used

Data from each individual was preprocessed from fastqs using SICILIAN with default parameters^26,51^. SICILIAN is a statistical method that can be applied to the BAM files by spliced aligners such as STAR to remove false positive junction calls, enabling unbiased discovery of unannotated junctions that can contribute to alternative splicing. The scRNA-Seq data sets were mapped using STAR version 2.7.5a in two-pass mode with default parameters. SpliZ scores were calculated using the SpliZ pipeline with default parameters^26,51^. Differential analysis was performed both based on tissue compartment (endothelial, epithelial, immune, and stromal), and independently based on cell type (defined by the tissue, compartment, and individual cell type, for example “lung immune macrophage”). The SpliZ was used to call genes as significant for all datasets except the full SS2 datasets, for which the SpliZVD was used because of the increased complexity of full-length transcript data. We used a p-value cutoff of 0.05 after Benjamini Hochberg correction. We define “effect size” for a gene to be the largest magnitude median SpliZ (or SpliZVD) value out of all cell types with calculable SpliZ for the gene. For between-cell-type-analysis, we use an effect size threshold of 0.5 (3.5 for SpliZVD) (Supplement), and require a difference of at least 0.5 within a single tissue and compartment for the gene to be called.

### FISH methods

Human Rectus Abdominus muscle biopsies from two donors were processed to single cell suspensions by a combination of manual and enzymatical dissociation (See TS paper). Single cell suspensions were stained with a combination of antibodies against CD45, CD31, THY1, and CD82, allowing for the isolation of immune cells (CD45+), endothelial cells (CD31+), mesenchymal cells (THY1+), and skeletal muscle satellite stem cells (CD82+). Due to the low number of immune cells present in the tissues, only the latter three cell types were stained. Cells were cytospun onto ECM coated 8-well chamber slides and fixed in 4% PFA. Cells were washed in PBS and prepared for RNA FISH by replacing the PBS to 100% ethanol. Cells were stained with custom probes according to the manufacturer’s protocol (Basescope Duplex Detection Reagent Kit (Advanced Cell Diagnostics = ACD). Briefly, cells were rehydrated and treated with Protease IV solution (1;15 dilution). Cells were subsequently stained with indicated Basescope probes for 2 hours in a hybridization oven set to 40 °C. Cells were then treated with amplification steps and imaged immediately after completion of the staining. As a control, human primary myoblasts were stained with the BaseScope probes and a Negative control probe. Images were captured with a Zeiss Axiofluor microscope with collected CCD camera and a 40x objective lens. The red dye fluoresces in the 555 channel, whereas the green dye shows as gray in the DIC channel. Images were quantified with Volocity software. One muscle sample was independently fixed in 10% neutral buffered formalin for 24 hours in preparation for BaseScope staining in cryosections. Tissue was dehydrated in 20% sucrose for 24 hours, washed in PBS, dried, embedded in OCT, and frozen in cooled isopentane. Sections of 10 µm were cut and dried in -20 °C for 1 hour and stored -80 °C until use. Tissue slides were removed from -80 °C and immediately washed with PBS to remove OCT, dried and baked in 60 °C for 30 min. Tissue slides were post-fixed in 4% PFA for 15 min and dehydrated by immersing slides in 50%, 70% and 100% ethanol for 5 min each. Tissue slides were then treated with hydrogen peroxide for 10 min and washed briefly with distilled water and subjected to target retrieval for 5 min, washed briefly with distilled water and in 100% ethanol. Tissue slides were treated with Protease IV solution in 40 °C for 10 min and washed twice with distilled water and hybridized with indicated BaseScope probes for 2 hours in 40 °C. Tissue slides were then treated with amplification steps. For dual FISH and IHC staining, tissue slides were immediately blocked in blocking buffer for 30 min (5% FBS, 1% BSA, 0.1% Triton-X100, 0.01% sodium azide in PBS) and stained with Pax7 antibody (1:100) in blocking buffer overnight in 4 °C. Tissue slides were washed with 0.1% tween-20 in PBS three times and then fluorescently conjugated secondary antibodies were added for an hour in room temperature. After three additional washes tissue slides were dried, mounted and imaged immediately.

De-identified human adult lung tissue was obtained from the Stanford Tissue Bank. The tissue was fixed in 10% neutral buffered formalin and embedded in paraffin. For the single-molecule in situ hybridization, 6um-thick paraffin sections were prepared and processed following the BaseScope Duplex Detection Reagent Kit (ACD) protocol, modified to use brown DAB chromogen in place of the usual green chromogen as the second color (custom protocol from ACD). Stained slides were visualized using an Olympus upright bright field microscope at 20x and 40x magnification. Cell types were identified by a pathologist based on cell morphology highlighted by the hematoxylin counterstain. Representative images of each cell type of interest were captured using an Olympus digital microscope color camera. Quantification was done by demarcating a polygonal image region containing multiple cells of homogeneous type and manually counting all the dots of each color within the region.

Proprietary probes (ACD) used for both human lung and muscle: BA-Hs-RPS24-tvc-1zz-st (targets 400-437 of NM_001026.5), BA-Hs-RPS24-tva-1zz-st (targets 399-437 of NM_033022.4); BA-Hs-MYL6-tv1-1zz-st (targets 469-505 of NM_021019.5), BA-Hs-MYL6-tv2-1zz-st (targets 436-480 of NM_079423.4).

### Single cell RT-PCR

Smart-seq2 preamplified cDNA of single cells from the Human Lung Cell Atlas project was used as starting templates^25^. The cells correspond to wells N14, A16, H14, B6, A3, A7, A13, A11, A8, D1, A12, A17, B12, J16, A21, P22, D23, A22, B22 of plate B002014; cell-type meta-data was taken from https://www.synapse.org/#!Synapse:syn21041850/wiki/60086. 1 µl of primary preamp was further preamplified in a 20 µl reaction (100 nM ISPCR primer = AAGCAGTGGTATCAACGCAGAGT, KAPA HiFi Fidelity mix; program: 95° 3’; 9 × [98° 20”; 67° 15”; 72° 4’]; 72° 5’), then diluted 8-fold with water. 2 µl of this secondary preamp was used as template in a 40 µl reaction (500 nM each of primers FL-RPS24ex4F1 = /6FAM/CAATGTTGGTGCTGGCAAAA and RPS24ex6R2 = GCAGCACCTTTACTCCTTCGG, New England Biolabs Phusion HF buffer, 200 nM dNTPs, 0.4 units Phusion DNA Polymerase; program: 98° 30”; 24 × [98° 10”; 60° 15”; 72° 20”]; 72° 5’). PCRs were diluted 1:100 and run on an ABI 3130xl Genetic Analyzer with GS500ROX standard; peaks were called by the Thermo Fisher Cloud Peak Scanner app, and presented as a pseudo-gel image using a custom Python script. For Sanger sequencing, secondary preamps were used in a similar PCR but with primers RPS24ex4F4 = AAGCAACGAAAGGAACGCAA and RPS24ex6R4 = CCACAGCTAACATCATTGCAG; the cleaned-up products were sequenced with the same primers. Oligonucleotide synthesis and capillary electrophoresis were done by Stanford PAN (Protein and Nucleic Acid Facility).

### Concordance analysis between technologies and donors

Concordance with Smart-seq2 was used as an extra test of the reproducibility of the method. Smart-seq2 and 10x datasets were subset to include only junctions and cell types shared in both to make the datasets as comparable as possible, and remove RNA measurements that could only be detected by Smart-seq2. Next, the SpliZ was calculated independently for both datasets as described for 10x. We then correlated the median SpliZ scores for matched genes and ontologies for genes called as significant by both technologies in the same individual. This resulted in a Pearson correlation of 0.439 between the two technologies for individual 1, and 0.769 for donor 2. We similarly subsetted both 10x datasets so that they each only included shared cell types and junctions, and then ran the SpliZ pipeline separately on each dataset, resulting in a Pearson correlation of 0.776 between the two datasets.

### K-means clustering of RPS24 and ATP5F1C

We first subset to only cells in the immune, epithelial, and stromal compartments with calculable SpliZ values for both RPS24 and ATP5F1C in the 10x data (9,712 cells in individual 1, 2,370 cells in individual 2). The endothelial compartment was not included because it had a small proportion of cells in both datasets (3.7% in individual 1, 4.5% in individual 2). K-means clustering was performed with sklearn in python with k=3 to separate all cells into three clusters based on their RPS24 and ATP5F1C SpliZ values. Each resulting cluster was assigned to one compartment such as to minimize classification error, and accuracy was calculated for each compartment based on these cluster assignments.

### LiftOver shared sites

We used the UCSC LiftOver tool (https://genome.ucsc.edu/cgi-bin/hgLiftOver) with the recommended settings to convert the the coordinates between human (hg38), mouse (mm10), and mouse lemur (Mmur3). To conserved variable splice sites between human, mouse, and mouse lemur, we analyzed only those junctions that have been successfully and uniquely converted by the LiftOver tool.

### Spermatogenesis analysis

To find genes with regulated splicing during sperm development, Spearman’s correlation was computed for each gene between the SpliZ and pseudotime values across the cells with computable SpliZ scores for that gene. We considered only genes with computable SpliZ in at least 100 cells and for each organism (human, mouse, mouse lemur), the genes with Spearman’s coefficient >0.1 and Bonferroni-corrected p-value <0.05 were selected as significantly splicing regulated genes. Only those genes that have names in all three organisms were considered for the conservation analysis.

## Tabula Sapiens Consortium

### Overall Project Direction and Coordination

Robert C Jones, Jim Karkanias, Mark Krasnow, Angela Oliveira Pisco, Stephen Quake, Julia Salzman, Nir Yosef

### Donor Recruitment

Bryan Bulthaup, Phillip Brown, Will Harper, Marisa Hemenez, Ravikumar Ponnusamy, Ahmad Salehi, Bhavani Sanagavarapu, Eileen Spallino

### Surgeons

Ksenia A. Aaron, Waldo Concepcion, James Gardner, Burnett Kelly, Nikole Neidlinger, Zifa Wang

### Logistical coordination

Sheela Crasta, Saroja Kolluru, Maurizio Morri, Angela Oliveira Pisco, Serena Y. Tan, Kyle J. Travaglini, Chenling Xu,

### Organ Processing

Marcela Alcántara-Hernández, Nicole Almanzar, Jane Antony, Benjamin Beyersdorf, Deviana Burhan, Lauren Byrnes, Kruti Calcuttawala, Mathew Carter, Charles K. F. Chan, Charles A. Chang, Alex Colville, Sheela Crasta, Rebecca Culver, Ivana Cvijović, Jessica D’Addabbo, Gaetano D’Amato, Camille Ezran, Francisco Galdos, Astrid Gillich, William R. Goodyer, Yan Hang, Alyssa Hayashi, Sahar Houshdaran, Xianxi Huang, Juan Irwin, SoRi Jang, Julia Vallve Juanico, Aaron M. Kershner, Soochi Kim, Bernhard Kiss, Saroja Kolluru, William Kong, Maya Kumar, Rebecca Leylek, Baoxiang Li, Shixuan Liu, Gabriel Loeb, Wan-Jin Lu, Shruti Mantri, Maxim Markovic, Patrick L. McAlpine, Ross Metzger, Antoine de Morree, Maurizio Morri, Karim Mrouj, Shravani Mukherjee, Tyler Muser, Patrick Neuhöfer, Thi Nguyen, Kimberly Perez, Ragini Phansalkar, Angela Oliveira Pisco, Nazan Puluca, Zhen Qi, Poorvi Rao, Hayley Raquer, Koki Sasagawa, Nicholas Schaum, Bronwyn Lane Scott, Bobak Seddighzadeh, Joe Segal, Sushmita Sen, Sean Spencer, Lea Steffes, Varun R. Subramaniam, Aditi Swarup, Michael Swift, Kyle J Travaglini, Will Van Treuren, Emily Trimm, Maggie Tsui, Sivakamasundari Vijayakumar, Kim Chi Vo, Sevahn K. Vorperian, Hannah Weinstein, Juliane Winkler, Timothy T.H. Wu, Jamie Xie, Andrea R.Yung, Yue Zhang,

### Sequencing

Angela M. Detweiler, Honey Mekonen, Norma Neff, Rene V. Sit, Michelle Tan, Jia Yan

### Histology

Gregory R. Bean, Gerald J. Berry, Vivek Charu, Erna Forgó, Brock A. Martin, Michael G. Ozawa, Oscar Silva, Serena Y. Tan, Pranathi Vemuri

### Computational Data Analysis

Shaked Afik, Rob Bierman, Olga Botvinnik, Ashley Byrne, Michelle Chen, Roozbeh Dehghannasiri, Angela Detweiler, Adam Gayoso, Qiqing Li, Gita Mahmoudabadi, Aaron McGeever, Antoine de Morree, Julia Olivieri, Madeline Park, Angela Oliveira Pisco, Neha Ravikumar, Julia Salzman, Geoff Stanley, Michael Swift, Michelle Tan, Weilun Tan, Sevahn Vorperian, Sheng Wang, Galen Xing, Chenling Xu, Nir Yosef

### Expert Cell Type Annotation

Marcela Alcántara-Hernández, Jane Antony, Charles A. Chang, Alex Colville, Sheela Crasta, Rebecca Culver, Camille Ezran, Astrid Gillich, Yan Hang, Juan Irwin, SoRi Jang, Aaron M. Kershner, William Kong, Rebecca Leylek, Gabriel Loeb, Ross Metzger, Antoine de Morree, Patrick Neuhöfer, Kimberly Perez, Ragini Phansalkar, Zhen Qi, Hayley Raquer, Bronwyn Lane Scott, Rahul Sinha, Hanbing Song, Sean Spencer, Aditi Swarup, Michael Swift, Kyle J. Travaglini, Jamie Xie

### Tissue Expert Principal Investigators

Steven E. Artandi, Philip Beachy, Michael F. Clarke, Linda Giudice, Franklin Huang, KC Huang, Juliana Idoyaga, Seung K Kim, Mark Krasnow, Christin Kuo, Patricia Nguyen, Stephen Quake, Thomas A. Rando, Kristy Red-Horse, Jeremy Reiter, Justin Sonnenburg, Bruce Wang, Albert Wu, Sean Wu, Tony Wyss-Coray

